# Polycomb safeguards imaginal disc specification through control of the Vestigial-Scalloped complex

**DOI:** 10.1101/2023.04.11.536444

**Authors:** Haley E. Brown, Brandon P. Weasner, Bonnie M. Weasner, Justin P. Kumar

## Abstract

A fundamental goal of developmental biology is to understand how cell and tissue fates are specified. The imaginal discs of *Drosophila* are excellent model systems for addressing this paradigm as their fate can be redirected when discs regenerate after injury or when key selector genes are mis-regulated. Here, we show that when *Polycomb* expression is reduced, the wing selector gene *vestigial* is ectopically activated. This leads to the inappropriate formation of the Vestigial-Scalloped complex which forces the eye to transform into a wing. We further demonstrate that disrupting this complex does not simply block wing formation or restore eye development. Instead, immunohistochemistry and high throughput genomic analysis show that the eye-antennal disc unexpectedly undergoes hyperplastic growth with multiple domains being organized into other imaginal discs and tissues. These findings provide insight into the complex developmental landscape that tissues must navigate before adopting their final fate.

**Summary Statement:** Here we describe a novel mechanism by which Pc promotes an eye fate during normal development and how the eye is reprogrammed into a wing in its absence.

## Introduction

A fundamental task of metazoans is to organize populations of cells into specialized tissues or organs. Cells first start out totipotent, transition through a pluripotency phase, and then enter a lineage-committed state (Ferrell, 2012; Riveiro and Brickman, 2020; Smith, 2017). This concept is enshrined in Waddington’s epigenetic landscape model of embryonic development which today forms the basis of modern developmental biology (Waddington, 1940, 1957). Since the model’s publication, numerous studies have shown that cells of one lineage can be converted into cells of another lineage through the simple expression of one or more transcription factors (Konstantinides and Desplan, 2020; Masserdotti et al., 2016; Takahashi and Yamanaka, 2015). One of the most striking examples is the forced expression of Pax6 induces the reprogramming of non-ocular tissues into retinal structures (Chow et al., 1999; Halder et al., 1995). Several transcription factors that induce cellular reprogramming interact directly with histone modifying proteins, thereby indicating that altering the chromatin landscape is important for transdetermination of tissue fate (Ang et al., 2011; Apostolou and Hochedlinger, 2013; Li et al., 2021; Mansour et al., 2012).

In this study we investigate the role that one epigenetic factor, Polycomb (Pc), plays in establishing the fate of the eye-antennal imaginal disc of the fruit fly, *Drosophila melanogaster*. Pc is a member of the Polycomb Group (PcG) of epigenetic repressors, and it is responsible for recognizing H3K27me3 (Cao et al., 2002; Czermin et al., 2002; Muller et al., 2002). Pc plays an important role in tissue specification as all thoracic and the first seven abdominal segments of null mutant embryos are transformed into the eighth abdominal segment (Lewis, 1978). Similarly, the adult T2 and T3 legs of a dominant *Pc* mutant are partially transformed into T1 legs (Lewis, 1947). Lastly, the frequency of leg-to-wing transdetermination events increases when PcG member levels are reduced (Lee et al., 2005).

Imaginal discs are excellent model systems for elucidating the mechanisms underlying cell and tissue specification in part because their fates can be dramatically altered by manipulations of key molecular determinants (Weasner and Kumar, 2022). For example, the fate of the eye is controlled by the retinal determination (RD) network of fourteen transcription factors (Kumar, 2010). In their absence, the developing eye field is transformed into head epidermal tissue (Hoge, 1915; Hunt, 1970; Mardon et al., 1994; Milani, 1941; Sved, 1986; Weasner and Kumar, 2013). Similarly, in the wing imaginal disc, the loss of several wing selector genes results in the complete loss of the wing blade (Ng et al., 1995; Sharma and Chopra, 1976; Stevens and Bryant, 1985; Williams et al., 1991). In contrast, the forced expression of eye and wing selector genes is sufficient to alter the fate of the targeted disc and provides additional insight into how tissue fate is controlled. For instance, forced expression of RD network members induces the formation of ectopic eyes in all imaginal discs (reviewed in Kumar, 2010). Similarly, ectopic activation of the wing selector gene vestigial (*vg*) transforms halteres into wings and induces wing outgrowths within the eye (Kim et al., 1996; Prasad et al., 2003; Simmonds et al., 1998).

Here, we report on the mechanisms by which the fate of the eye-antennal disc is specified. When this disc is fragmented. regenerating cells along the wounded edge are often reprogrammed and give rise to wing tissue (Hadorn, 1968, 1978). Reprogramming of the eye into a wing also occurs in response to several genetic perturbations (Edwards and Gardner, 1966; Goldschmidt and Lederman-Klein, 1958; Katsuyama et al., 2005; Kobel, 1968; Kurata et al., 2000; Masuko et al., 2018; Ouweneel, 1970; Postlethwait, 1974; Simmonds et al., 1998). Interestingly, there are no recorded instances of the eye being transformed into any other imaginal disc. This contrasts with the adjacent antenna, which can be reprogrammed to develop into wing, leg, labial, and genital discs (Hadorn, 1978; Weasner and Kumar, 2022). We set out to understand why the eye field is specifically reprogrammed into a wing disc rather than one of the other imaginal discs.

In a previous study we discovered that a reduction in *Pc* expression reprograms the eye into a wing (Zhu et al., 2018). Here we show that reducing *Pc* expression results in the ectopic activation of *vg* within the eye field. The repression of *vg* expression by Pc during normal eye development is likely to be direct as functional Polycomb Response Elements (PREs) are present within the *vg* locus (Ahmad and Spens, 2019; Herzog et al., 2014; Okulski et al., 2011). In the developing wing Vg interacts with the Scalloped (Sd) DNA-binding protein to promote the fate of the wing (Delanoue et al., 2004; Halder and Carroll, 2001; Halder et al., 1998; Kim et al., 1996; Klein and Arias, 1998; Pimmett et al., 2017; Simmonds et al., 1998). If this complex is disrupted then the adult wing is severely reduced in size or eliminated if the Vg-Sd complex is disrupted (Bownes et al., 1981; Gruneberg, 1929; Lindsley and Grell, 1968; Williams and Bell, 1988). In contrast, since *vg* is not normally expressed in the developing eye, Sd interacts with the Yorki transcriptional activator instead. The Yki-Sd complex goes on to regulate the final size of the compound eye (Koontz et al., 2013; Meserve and Duronio, 2015). Our results suggest that the eye field is prevented from developing into a wing by Pc mediated repression of the *vg* locus. In the absence of this repression, ectopic activation of Vg within the eye field allows for the inappropriate formation of the Vg-Sd complex and the reprogramming of the eye into a wing. Thus, our findings suggest that selective regulation of Sd binding partners by PcG proteins guides the eye primordium along the path towards its final fate.

We made a second surprising discovery when we reduced *Pc* expression in *vg* and *sd* loss-of-function mutant backgrounds to a way to prevent the Vg-Sd complex from forming within the eye-antennal disc. Instead of the disc simply reverting to its normal state, it instead undergoes massive hyperplastic growth. This is reminiscent of the immense expansion seen when certain Polycomb Group (PcG) factors are misregulated (Beira et al., 2018; Bunker et al., 2015; Classen et al., 2009; Loubiere et al., 2016; Martinez et al., 2009) or when epithelial polarity is lost (Bunker et al., 2015; Enomoto et al., 2021; Pagliarini and Xu, 2003; Wu et al., 2010) in the eye-antennal disc. The explosive growth that we observe is consistent with the role of Sd as a default repressor of tissue growth (Koontz et al., 2013). Unexpectedly, histological analysis of the mutant tissue indicates that multiple domains of the overgrown discs take on the fate of other imaginal discs. RNA sequencing (RNA-seq) confirms the presence of several exogenous imaginal discs as well as several other distinct tissue types. Our results bring to mind the diverse collection of cell types that are found within mammalian tumors (Dagogo-Jack and Shaw, 2018; Genovese et al., 2019; Ju, 2021; Li et al., 2022; McGranahan and Swanton, 2017; Tammela and Sage, 2020). As such, information gleaned from hyperplastic imaginal discs, may also provide insight into mechanisms that give rise to intratumor heterogeneity. In total, our results elucidate the complex developmental landscape that tissues must navigate on the way to adopting final fates.

## Results

### The loss of *Pc* results in an eye-to-wing transformation

A prior study showed that, upon reduction of *Pc,* the dorsal eye field is transformed into a wing imaginal disc. These discs show ectopic activation of the *Antp* Hox gene and the *vg* wing selector gene as well as the loss of the RD network genes *ey* and *eya* (Zhu et al., 2018). The current model is that, during normal eye-antennal disc development, Pc promotes eye fate by epigenetically repressing the expression of wing fate genes. Removal of this inherent repression results in the inappropriate activation of wing selector genes and the transformation of the eye into a wing. Since simultaneous reductions in *Pc* and *Antp* levels results in dramatically fewer instances of eye-to-wing transformations, it was proposed that *Antp* plays a central role in this fate decision. However, the eye-to-wing transformation is not observed in most *Antp* gain-of-function mutants (Scott et al., 1983) or when *Antp* is forcibly expressed in the developing eye (Jorgensen and Garber, 1987; Kurata et al., 2000; Schneuwly et al., 1987b). As such, the mechanism by which Pc maintains the fate of the eye remains unknown.

To investigate the molecular mechanism underlying the transformation of an eye into a wing, we used RNA interference (RNAi) to individually knock down 18 of 20 PcG members in the eye-antennal disc (Figure 1, Table S1). PcG factors are ubiquitously expressed across different tissues (Buchenau et al., 1998; DeCamillis and Brock, 1994; Paro and Zink, 1993) and consistent with these studies, our RNA-seq data (Table S2) shows that transcripts of all PcG members, and their known binding partners, are expressed within wild-type eye-antennal and wing discs as well as *Pc* knockdown eye-antennal discs (Figure S1A-D). Therefore, knocking down PcG members in the eye-antennal disc should enable us to elucidate the function of these factors in eye development. Since we focus on the effects of knocking down *Pc*, we validated the *Pc* RNAi line by showing that *Pc* transcript levels are reduced in knockdown discs compared to wild-type eye-antennal discs (Figure S1E). Using a Pc specific antibody we have also shown the RNAi line to be effective in knocking down *Pc* expression in the eye field (Zhu et al., 2018).

**Figure 1.**
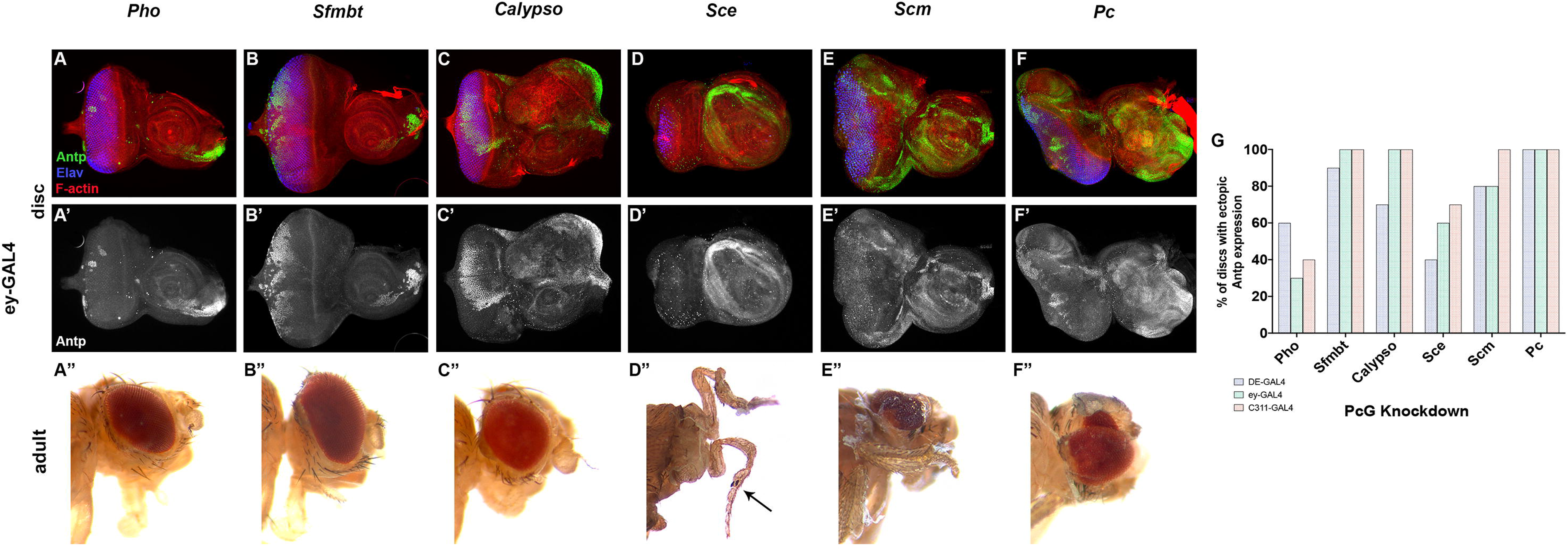
Antp is ectopically expressed when some PcG members are knocked down. (A-F) Third-instar eye-antennal imaginal discs stained with anti-Antp (green), anti-Elav (blue), and phalloidin (red). (A’-F’) Antp channel alone from the same eye-antennal discs. (A”-F”) Adults lacking either *Pho* (A”), *Sfmbt* (B”), or *Calypso* (C”) can eclose and maintain mostly wild-type morphology. However, knocking down either *Sce* (D”), *Scm* (E”), or *Pc* (F”) is pharate lethal. Arrow (D”) marks the presence of a sex comb indicating that this is a T1 leg. (G) Bar graph represents percent of PcG knockdown discs that show ectopic *Antp* expression with three different GAL4 drivers (*DE-GAL4*: blue; *ey-GAL4*: green; *c311-GAL4*: orange).

Since *Antp* is robustly expressed in *Pc* knockdown eye-antennal discs and required for wing formation (Fang et al., 2022; Paul et al., 2021; Zhu et al., 2018), we first screened for ectopic activation of *Antp* and a transformation of the eye into a wing. Surprisingly, while we detected *Antp* expression when knocking down five other PcG members (*pleiohomeotic* (*pho)*, *Scm-related gene containing four MBT domains* (*Sfmbt)*, *Sex combs extra* (*Sce)*, *Sex comb on midleg* (*Scm)* and *Calypso*), the expression levels and pattern of *Antp* in these discs differs from the *Pc* knockdowns and the eye does not transform into a wing. In *pho*, *Sfmbt*, and *Calypso* knockdown discs, ectopic *Antp* expression is largely restricted to the peripodial epithelium (Figure 1A-C). As such, the corresponding adults have bristle, head capsule, and antennal defects but relatively wild-type compound eyes (Figure 1A”-C”). Reductions of *Sce* results in the expression of ectopic *Antp* just within the antennal segment (Figure 1D). This differs drastically from that of *Pc* knockdown discs, where *Antp* is expressed throughout the antennal field and the ectopic wing tissue (Figure 1F). As such, this may explain why the reduction of *Sce* induces a full antenna-to-leg transformation (Figure 1D”) but not the eye-to-wing transformation seen in *Pc* knockdown discs (Figure 1F”). Interestingly, in contrast to dominant gain-of-function *Antp* mutants in which the antenna is transformed into the second thoracic T2 leg (le Calvez, 1948a, b, c; Yu, 1949), reductions in *Sce* transform the antenna into a sex comb bearing T1 leg (Figure 1D”, arrow). Lastly, while knocking down *Scm* results in ectopic *Antp* expression within the eye and antenna, only the fate of the antenna is altered (Figure 1E-E”). These five ectopic *Antp* expression patterns are consistently observed when PcG factors are knocked down with three different GAL4 drivers: *DE-GAL4* (dorsal eye field), *ey-GAL4* (throughout the eye field), and *c311-GAL4* (peripodial epithelium) (Figure 1G).

Since ectopic *Antp* expression in multiple PcG member knockdowns is not associated with eye-to-wing transformation, these results suggest that, in contrast to the earlier published model, the eye-to-wing transformation is caused, not by the de-repression of *Antp*, but rather by the potential de-repression of other wing selector genes. This is more in line with studies suggesting that *Antp* plays only a minor role in forewing development (Carroll et al., 1995; Tomoyasu et al., 2005). Our findings build upon the idea that eye fate is normally driven by Pax6-mediated activation of the retinal determination gene regulatory network (Gehring and Ikeo, 1999; Kumar, 2010) and maintained by *Pc*-dependent repression of non-Hox wing selector genes.

### HOX gene misregulation disrupts the programmed eye fate

Since Pc functions to establish the expression pattern of Hox genes during embryonic development (Janody et al., 2004; Lewis, 1978; Ringrose and Paro, 2007; Schwartz et al., 2006; Struhl and Akam, 1985; Tolhuis et al., 2006) and to maintain imaginal disc fate (Katsuyama and Paro, 2011; Lee et al., 2005), the perturbation of PcG repression might be expected to result in the ectopic expression of these genes within the eye field. Transcript counts from RNA-seq analysis of wild-type and *Pc* knockdown eye-antennal discs indicate that the eight HOX genes can be divided into two groups when *Pc* is knocked down (Figure 2A”-H”). The first group includes *Sex combs reduced* (*Scr*), *Antp*, *Ultrabithorax* (*Ubx*), and *Abdominal-B* (*Abd-B*) – these genes are normally silenced within the eye-antennal disc, but their expression is elevated within the disc upon *Pc* knockdown (Figure 2A’’-D’’). The second group is comprised of *labial* (*lab*), *proboscipedia (pb)*, *Deformed* (*Dfd*), and *abdominal-A* (*abd-A*). The expression of these four Hox genes does not change significantly in *Pc* knockdown discs (Figure 2E’’-H’’). Of these, *pb* and *Dfd* are normally expressed within the developing head epidermis and peripodial epithelium but not the eye (Aplin and Kaufman, 1997; Diederich et al., 1991).

**Figure 2.**
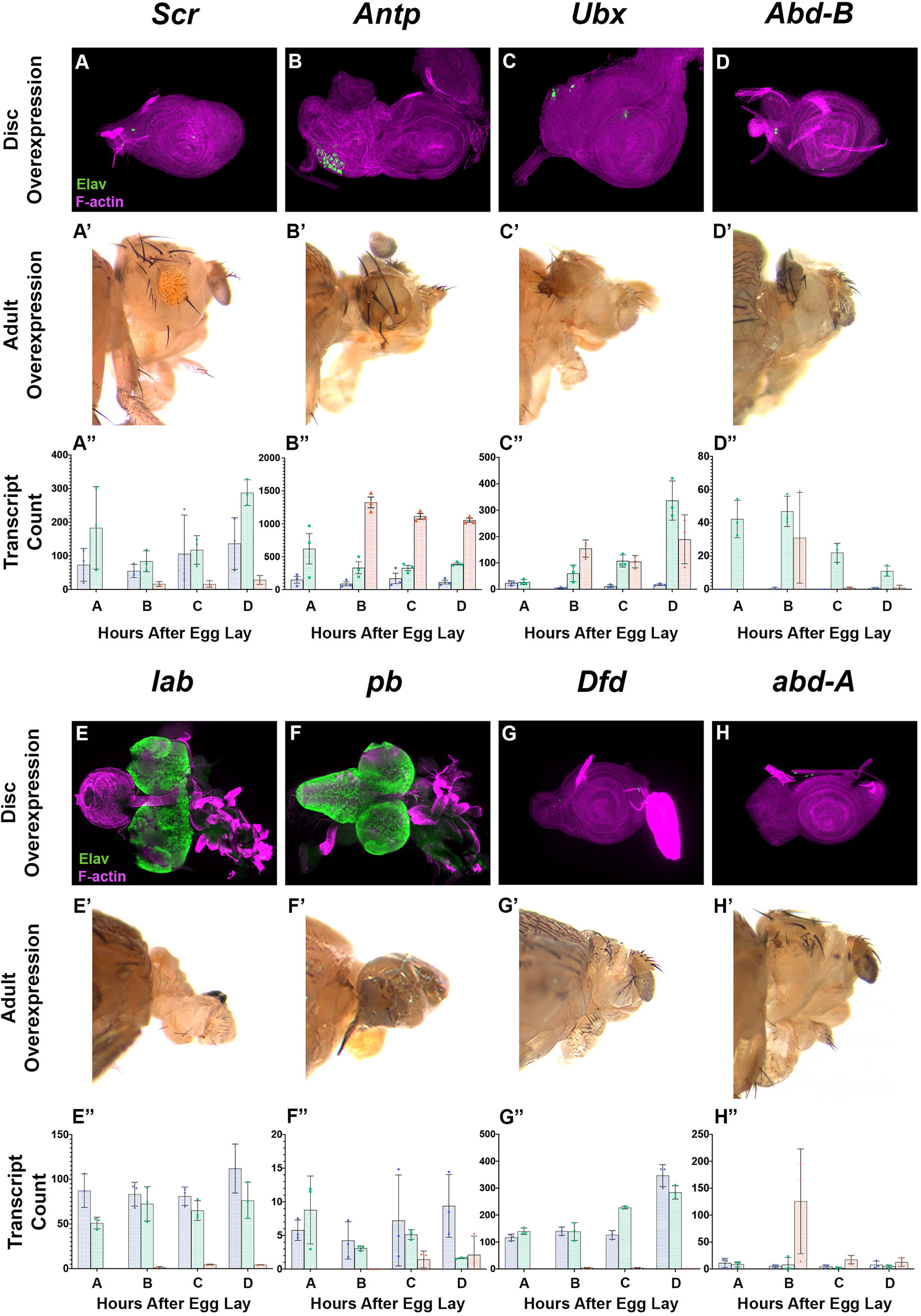
Overexpression of HOX genes disrupts eye development. (A-D) Hox genes with elevated transcript levels in *Pc* knockdown discs compared to wild-type eye-antennal discs (*Scr*, *Antp*, *Ubx*, and *Abd-B*). (E-H) HOX genes with relatively unchanged transcript levels in *Pc* knockdown discs compared to wild-type eye-antennal discs (*lab*, *pb*, *Dfd*, and *abd-A*). (A-H) Third instar eye-antennal imaginal discs stained with anti-Elav (green) and phalloidin (magenta). Whole mounted disc-brain complexes are shown for *ey*>*lab* (E) and *ey*>*pb* (F). (A’-H’) Eclosed adult flies are eyeless and missing head capsule (G’, H’) and have reduced number of ocelli (B’-D’). Pharate lethal flies are headless (E’, F’). (A”-H”) Bar graphs represent DESeq2 normalized transcript counts of three replicates for the wild-type eye-antennal disc (blue), *Pc* knockdown disc (green), and wild-type wing disc (orange). Comparisons comprise third instar larval development; where A: 84hr WT EAD v. 96hr *PcKD*, B: 96hr WT v. 108hr *PcKD*, C: 108hr WT v. 120hr *PcKD*, and D: 120hr WT v. 144hr *PcKD* (error bars represent standard deviation).

To determine the impact that ectopic expression of the Hox genes might play in the *Pc*-dependent eye-to-wing transformation, we forcibly expressed each of the eight Hox genes within the developing eye field using *ey-GAL4* (Figure 2, Table S1). Consistent with prior reports, the overexpression of *Antp* alone results in either eyeless (83%) or headless (17%) flies but no eye-to-wing transformations (Figure 2B,B’). So, while *Antp* may contribute to the establishment of this transdetermination event, it alone is not sufficient to do so, consistent with our observations in other PcG knockdowns. Like the forced expression of *Antp*, the overexpression of several remaining Hox genes severely reduced the size of the eye field and thus resulted in predominantly eyeless adults (Figure 2A-D, G-H). The effect of ectopic *Scr* on the developing eye is slightly milder with 52% of adults having small eyes (Figure 2A,A’) while the rest are eyeless. Overexpression of *pb* and *lab* eliminated the eye-antennal disc completely (Figure 2E-F) and resulted in headless pharate adults (Figure 2E’-F’). These results show that Hox gene overexpression in the eye field severely impairs eye, ocellar, and head fate. Antp and Pb proteins, when ectopically expressed within the eye-antennal disc, inhibit eye development by physically binding to Ey and preventing it from binding to its target enhancers (Benassayag et al., 2003; Plaza et al., 2008; Plaza et al., 2001). Eye development may be disrupted by a similar mechanism when the other Hox genes are upregulated. Our hypothesis is that inappropriate levels of Hox genes, which result from the loss of *Pc*, destroy the pre-programmed eye fate. Under these circumstances, the eye field is primed for other selector gene(s) to reprogram the eye into a wing.

### Bulk RNA-seq analysis identifies candidates that play a role in the eye-to-wing transformation

To identify putative selector genes underlying the eye-to-wing transdetermination event, we used bulk RNA-seq to compare the transcriptome profiles of *Pc* knockdown eye-antennal discs (*ey>Pc RNAi*) to wild-type (WT) eye-antennal and wing discs throughout third larval instar development (96, 108, 120 and 144hrs after egg lay (AEL)) (Figure S2, Table S2). When *Pc* is knocked down, eye discs are developmentally delayed by approximately 24 hours – the larvae pupate at 144hrs instead of 120hrs AEL. This delay is consistent throughout the developmental window described above, as hierarchical clustering analysis shows that the transcriptome of each *Pc* knockdown disc roughly clusters with that of the 24hr earlier WT eye-antennal disc (Figure S2B). We therefore compared the transcriptome of *Pc* knockdown discs to WT eye-antennal and wing discs throughout ‘early’ (108hr v. 96hr AEL), ‘mid’ (120hr v. 108hr AEL), and ‘late’ (144hr v. 120hr AEL) third instar development (Figure S3A-C). Differential expression analysis (log2 fold change, adjusted p<0.05) of these three developmental windows identified 516 early genes, 230 mid genes, and 233 late genes with elevated expression in both *Pc* knockdown eye-antennal discs and WT wing discs when compared to WT eye-antennal discs (Figure 3A-D). We then compared these candidate lists to each other to determine which genes are potentially underlying the *Pc*-dependent eye-to-wing transformation. While numerous genes are specific to early, mid, and late development, there are also several candidates which are elevated during multiple stages (Figure 3E-F, Table S3). Enriched Gene Ontology (GO) analysis shows that early genes are mainly important for lipid metabolism and biosynthetic processes (Figure S3D). While we found no significant enrichment clusters in the mid or late candidates, the category shared by early and mid-stage genes showed enrichment for fatty acid biosynthesis, metabolism, and translation elongation (Figure S3E).

**Figure 3.**
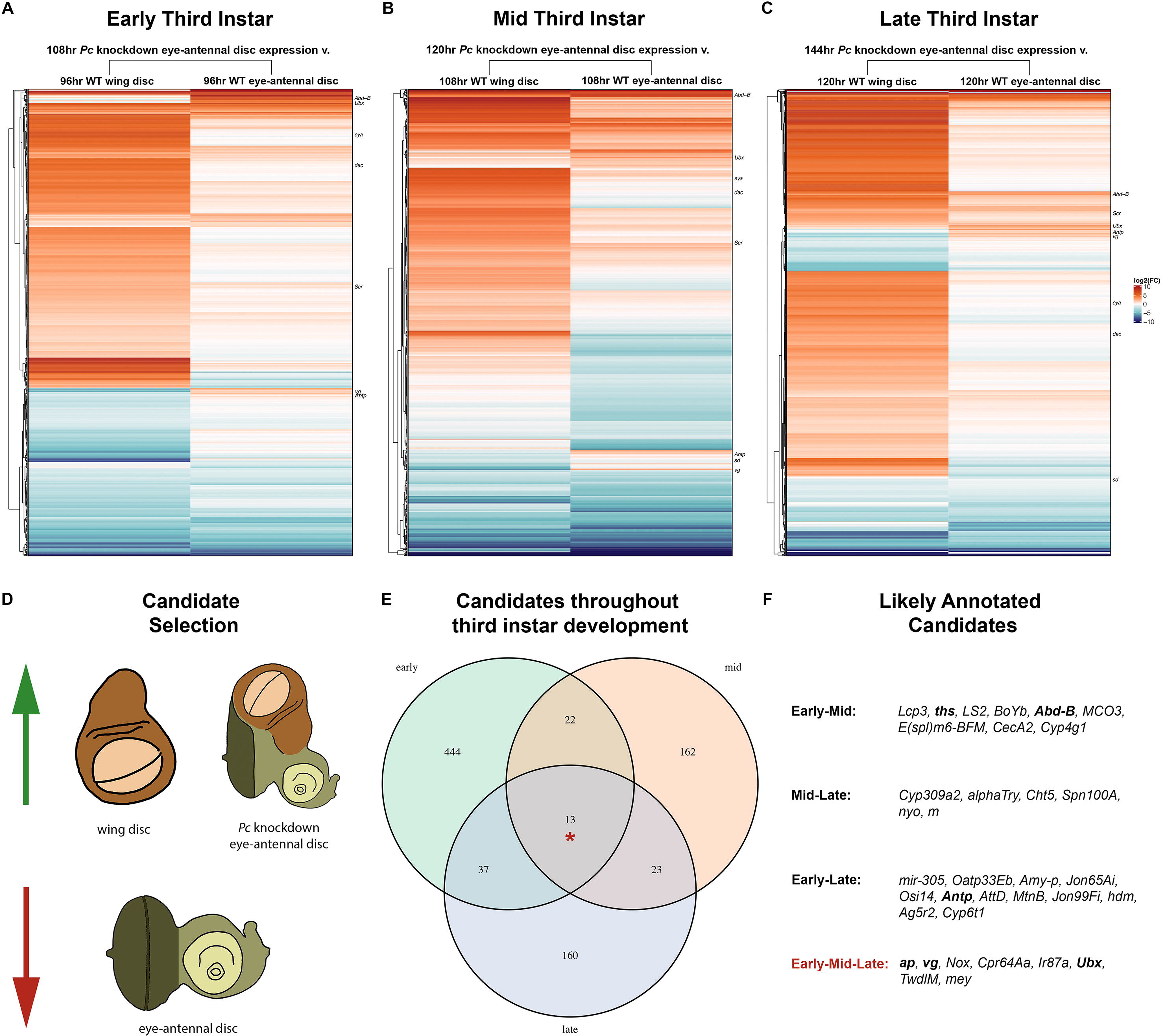
Differential expression of genes in response to *Pc* reduction. (A-C) Heatmap of log2 transformed differential transcript expression between *Pc* knockdown eye-antennal discs and wild-type (WT) wing discs (left) or eye-antennal discs (right) throughout early (A, 108hr v. 96hr), mid (B, 120hr v. 108hr), and late (C, 144hr v. 120hr) third instar development. Known eye genes, wing genes, and Hox genes are listed to the right. (D) Schematic representing selection method for potential candidates behind the eye-to-wing transformation. (E) Venn diagram showing the overlap of candidate genes throughout early (green, n = 516), mid (orange, n = 220), and late (blue, n = 233) third instar development. Asterisk denotes candidate genes that are differentially expressed throughout the entire tested developmental window. (F) List of annotated candidates in each of the overlapping categories. Bold face represents genes with available UAS overexpression lines. Remaining genes for each category are listed in Supplemental Table 3.

Our RNA-seq data shows that *Antp* is only enriched in *Pc* knockdown discs during early and late third instar development (Figure 3F). This provides additional support for our model that *Antp* may be required, but is not sufficient, for the eye-to-wing transformation. We identified other wing selector genes, such as *nubbin* (*nub*), being enriched only at late developmental stages. These genes may be pivotal for continuing the transdetermination of the eye into a wing but are unlikely to be responsible for initiating the switch in fate. We ectopically expressed genes with available UAS lines (Figure 3F, boldface) with *ey-GAL4* and found that the wing genes *nub* and *wingless (wg)* are insufficient to transform the eye into a wing (Table S1). As genes that are upregulated at just one or two developmental stages cannot induce the eye-to-wing transformation, we hypothesized that to initiate transdetermination, and maintain the push towards a novel fate, selector genes should be consistently enriched in *Pc* knockdown discs at all developmental stages. Thirteen candidates comprise this category and notably include the Hox gene *Ubx* and two wing selector genes – *vg* and *apterous* (*ap*). We further hypothesized that reductions in *Pc* within the eye-antennal disc should lower the levels of the PcG-dependent repressive mark H3K27me3 at these loci and allow for these genes to be ectopically expressed. These factors would then be able to inappropriately activate the downstream wing gene regulatory network and force a transformation of the eye into a wing.

To test this hypothesis, we performed Cleavage Under Targets & Release Using Nuclease (CUT&RUN) on whole imaginal discs and determined the profile of PcG-specific marks (H3K27me3 and H2AK119Ub) in *Pc* knockdown eye-antennal discs as well as both wild-type eye-antennal and wing discs (Figure 4, S4). We find that genome-wide H3K27me3 peaks in *Pc* knockdown eye-antennal discs resemble that of wild-type wing discs more closely than wild-type eye-antennal discs (Figure 4A). However, the H2AK119Ub profiles of *Pc* knockdown discs appear more similar to wild type eye-antennal discs (Figure 4B). This is not surprising as the role that ubiquitination plays in PcG repression is not fully understood (Blackledge et al., 2014; Cohen et al., 2020; Cooper et al., 2014; Kahn et al., 2016; Kalb et al., 2014; McGinty et al., 2014; Pengelly et al., 2015; Tamburri et al., 2020). An examination of peak profiles within candidate genes expressed throughout third larval instar development (Figure 4C) shows that the 13 genes fall into three categories: (1) elevated transcript levels and diminished H3K27me3, (2) elevated transcript levels but minimally changed H3K27me3, and (3) candidates with generally low transcript counts (>5). As 8 of the 13 genes show elevated transcript levels and diminished levels of H3K27me3 in *Pc* knockdown discs compared to wild-type eye-antennal discs (category 1), this supports our hypothesis that reduction of *Pc* expression prevents either the maintenance or propagation of H3K27me3 and this ultimately allows candidates to be ectopically expressed and alter the fate of the eye-antennal disc.

**Figure 4.**
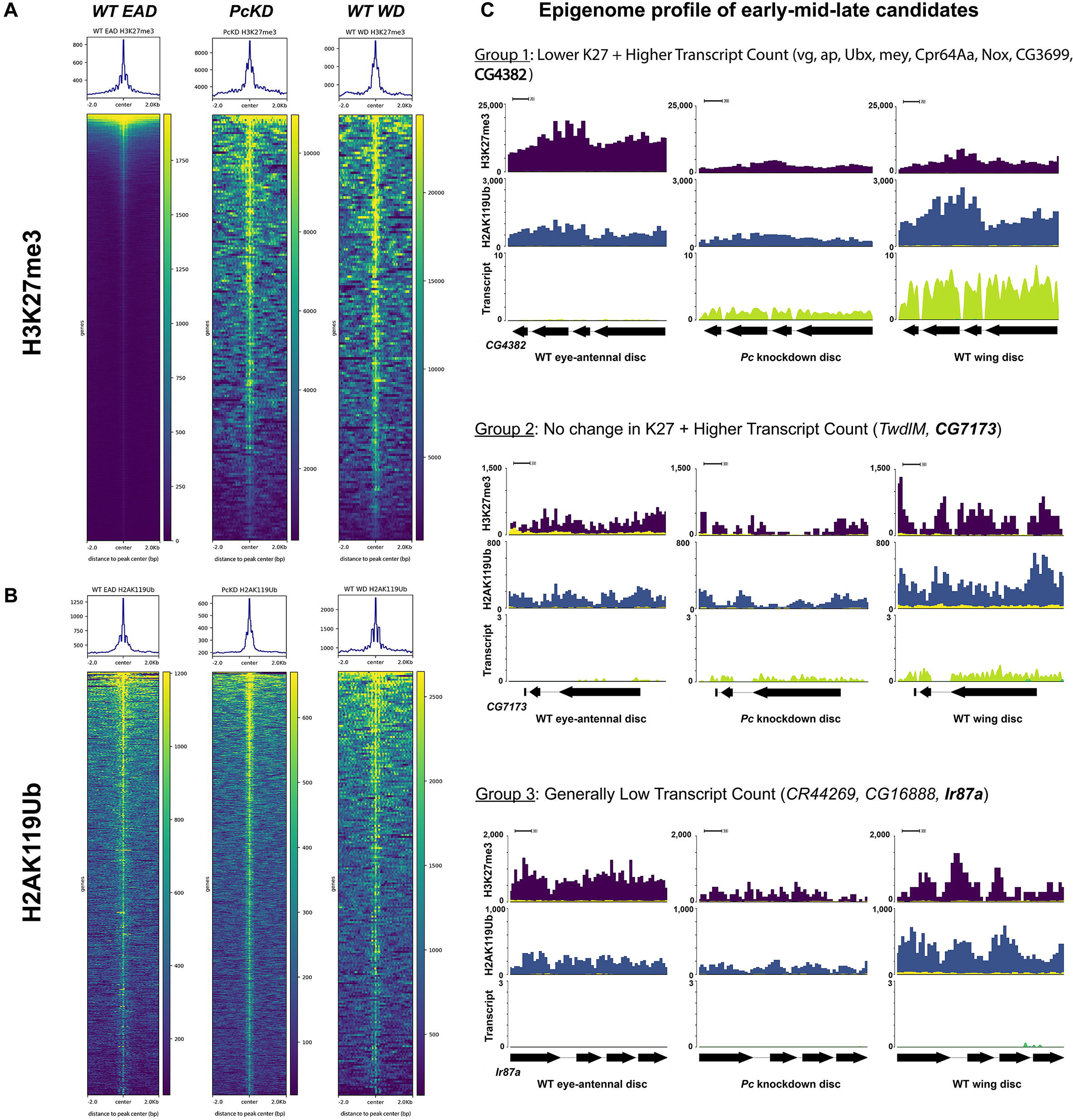
*Pc* reduction results in H3K27me3 profiles that resemble wing discs. (A) H3K27me3 and (B) H2AK119Ub CUT&RUN heatmaps of wild-type eye-antennal discs (EAD, left), *Pc* knockdown eye-antennal discs (PcKD, center), and wild-type wing discs (WD, right). (C) CUT&RUN H3K27me3 (indigo), H2AK119Ub (blue), and transcript (green) peaks at the *CG4382* (Group 1), *CG7173* (Group 2), and *Ir87a* (Group 3) loci of each genotype. The corresponding histone mark IgG control track is overlayed in yellow.

### The wing selector gene *vestigial* alters the fate of the eye-antennal disc

As described above, several wing genes were identified as candidates for forcing the eye-to-wing transformation. The most promising candidate is *vg*, as forced expression of it alone in the eye field induces an eye-to-wing transformation similar to what we observe in *Pc* knockdown discs (Kim et al., 1996; Simmonds et al., 1998) (Figure 5A,D). This ability remains unique to *vg* as overexpressing other wing selector genes (including *ap*, *nub*, and *wg*) did not induce any wing-like outgrowths (Table S1). During normal development, *vg* is expressed in a broad stripe within the wing pouch but is entirely absent from the eye-antennal disc (Figure 5B,C) (Kim et al., 1996; Williams et al., 1991; Zhu et al., 2018). In transformed *Pc* knockdown discs, *vg* transcripts are differentially expressed throughout third instar development and Vg protein is now localized exclusively to the pouch region of the ectopic wing (Figure 5E,F). The presence of two functional PREs within the *vg* locus (Ahmad and Spens, 2019; Herzog et al., 2014; Okulski et al., 2011; Ringrose et al., 2003; Srinivasan and Mishra, 2020) suggests that it is repressed by PcG proteins during normal development of the eye-antennal disc (Figure 5G, asterisks).

**Figure 5.**
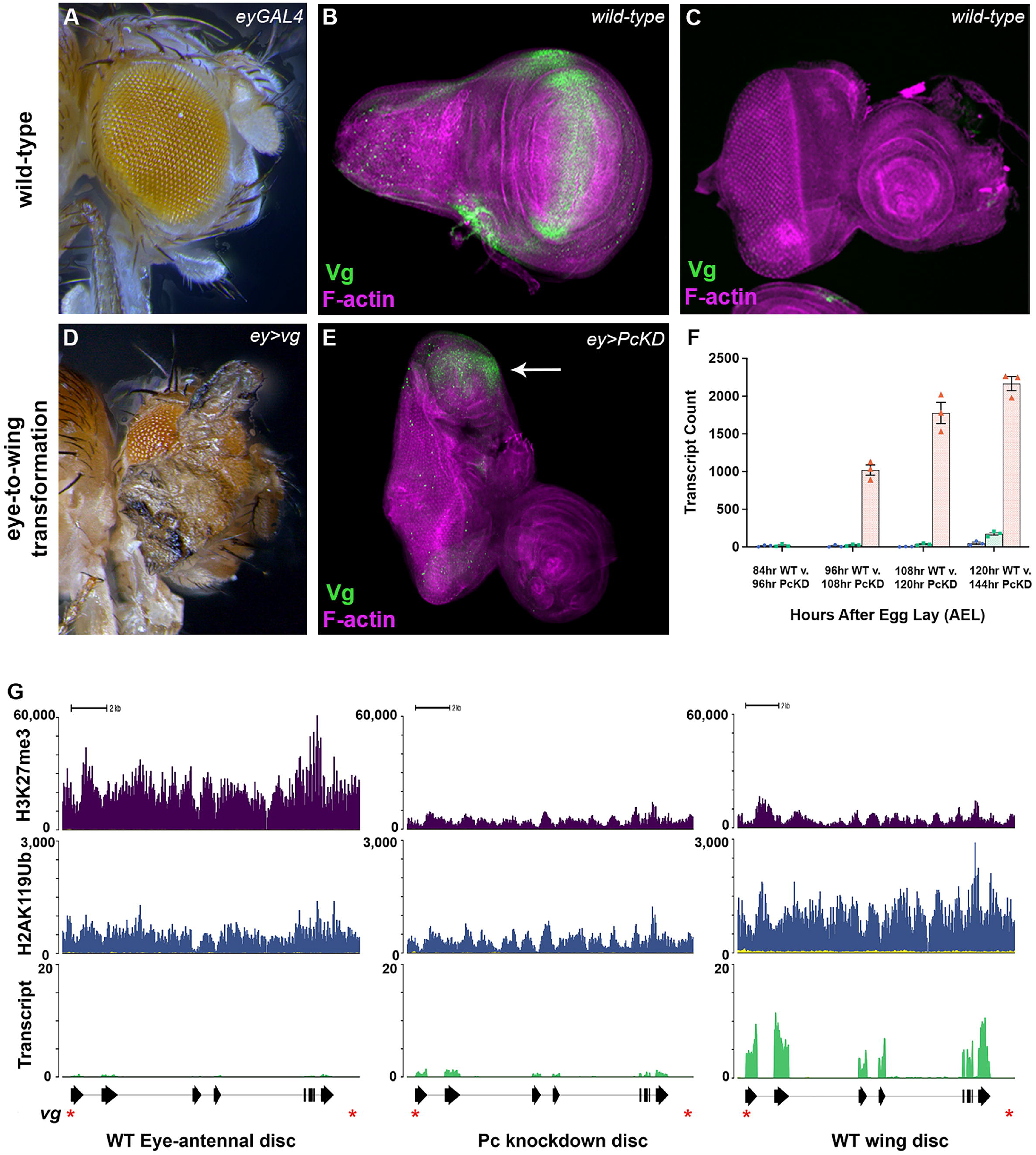
Vestigial is a candidate for the Pc eye-to-wing transformation. (A) Wild-type adult head. (B-C) Wild-type third instar wing disc (B) and eye-antennal disc (C) stained with anti-Vg (green) and phalloidin (magenta). (D) Overexpression of *vg* in the eye-antennal disc is sufficient to drive an eye-to-wing transformation. (E) *ey>Pc* RNAi transformed discs stained with anti-Vg (green) and phalloidin (magenta). Vg is ectopically activated in the pouch of the transformed disc (arrow). (F) Bar graphs represent DESeq2 normalized transcript counts of three replicates for the wild-type eye-antennal disc (blue), *Pc* knockdown disc (green), and wild-type wing disc (orange). Error bars represent standard deviation. (G) CUT&RUN H3K27me3 (top), H2AK119Ub (middle), and forward transcript (bottom) peaks at the *vg* locus in the wild-type eye-antennal disc, *Pc* knockdown disc, and wild-type wing disc. The corresponding histone mark IgG control track is overlayed in yellow. Asterisks denote the two PREs identified by Ahmad and Spens, 2019.

Our CUT&RUN analysis (Figure 5G) further supports this hypothesis as we found high levels of H3K27me3 at the *vg* locus in the eye disc (where *vg* would normally be repressed). Conversely, in the wing disc (where *vg* is expressed across the wing pouch) the *vg* locus is decorated with significantly lower levels of H3K27me3. The continued presence of H3K27me3 peaks (albeit at lower levels) is expected since we analyzed whole wing discs that contains a mixture of *vg* expressing and non-expressing cells. In *Pc* knockdown eye-antennal discs H3K27me3 levels at the *vg* locus are drastically reduced and appear similar to wing discs. Although the chromatin profile H2AK119Ub is also altered in *Pc* knockdown discs it does not truly resemble the profile of either eye or wing discs. These results demonstrate that when *Pc* levels are reduced, the repressive chromatin landscape of *vg* is diminished, and this results in it being ectopically expressed within the transdetermining eye-antennal disc.

### Scalloped is required for Vg-dependent transformation of the eye into a wing

While ectopic Vg in the eye (Kim et al., 1996; Simmonds et al., 1998) can yield wing outgrowths, the question of how Vg is able to convert this tissue remains. In the wing disc, its binding partner, Sd, is enriched within a similar stripe across the pouch. The Vg-Sd complex promotes wing fate by activating downstream wing fate genes (Garg and Bell, 2010; Halder and Carroll, 2001; Halder et al., 1998). Sd also promotes growth of the wing pouch by forming a complex with the Yorkie (Yki) transcriptional activator (Koontz et al., 2013; Wu et al., 2008). In the developing eye (where Vg is absent), Sd is present at low levels throughout the disc and promotes its growth via the Yki-Sd complex (Campbell et al., 1992; Guss et al., 2013; Koontz et al., 2013; Wu et al., 2008). A possible mechanism underlying the eye-to-wing transformation is that Vg, which is now present in the eye because of knocking down *Pc*, competes with Yki for binding to Sd. This could recreate the genetic conditions that are present within the wing disc and, as a result, the eye field would be reprogrammed and develop into a wing. To determine whether Sd is required to induce the formation of wing outgrowths that are caused by ectopic Vg, we knocked down *sd* in animals that are overexpressing *vg* (Figure 6A,C and A’,C’). As expected the adult eye no longer transforms into wing tissue and instead appears like that of *sd* knock down animals (Figure 6B,B’). We conclude that the eye-to-wing transformation seen in *Pc* knockdown animals is due to ectopic Vg taking advantage of Sd that is already present in the eye-antennal disc. The Vg-Sd complex forces the eye field to transdetermine into a wing by then initiating activation of the downstream wing gene regulatory network.

**Figure 6.**
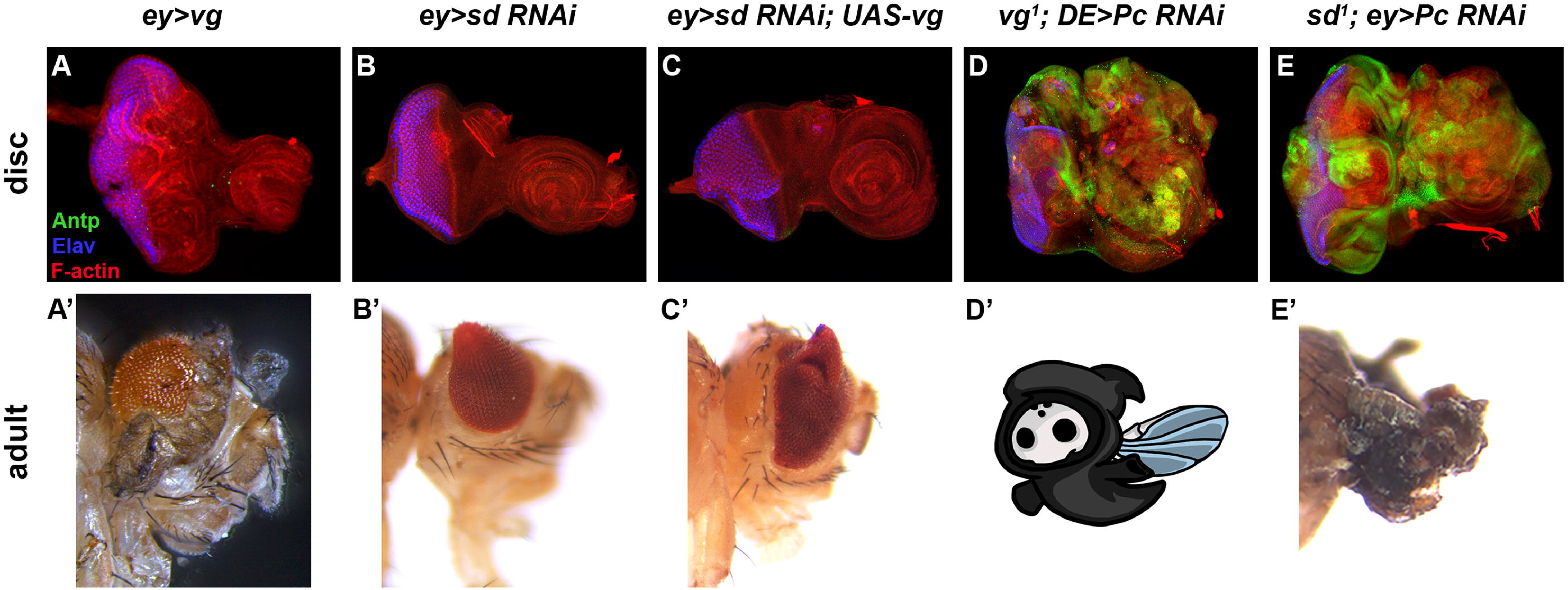
The Pc-dependent eye-to-wing transformation relies on the Vg-Sd complex. (A-E) Third-instar eye-antennal imaginal discs stained with anti-Antp (green), anti-Elav (blue), and phalloidin (red). (A’-E’) Eclosed or pharate lethal adult heads of the corresponding genotype. (A) Overexpression of *vg* disrupts photoreceptor development and results in disc folds in the anterior eye field. (A’) Eclosed adults develop wing outgrowths on every axis of the eye. (B-B’) Knockdown of *sd* alone results in malformations in the morphology of the anterior eye field (B) and adults with abnormal, cone-shaped eye outgrowths (B’). (C-C’) When *vg* is overexpressed while *sd* is knocked down, the phenotype mimics *sd* knockdown animals (B’) rather than *vg* overexpression animals (A’). (D-E) Knocking down *Pc* in a *vg^1^* (D) or *sd^1^* (E) mutant background disrupts the eye-to-wing transformation and instead leads to hyperproliferative discs. (D’-E’) The combined loss of *vg* and *Pc* (D’) is pupal lethal, while the combined loss of *sd* and *Pc* (E’) is pharate lethal and results in flies with diminished, ‘tumorous’ heads.

### Ablating the Vg-Sd complex in *Pc* knockdown discs induces multiple new imaginal disc fates

To further test whether the formation of the Vg-Sd complex drives the *Pc*-dependent eye-to-wing transformation, we genetically disrupted the complex by knocking down *Pc* in *vg^1^* or *sd^1^*loss-of-function mutant backgrounds. We expected that loss of either gene would prevent the eye from being transformed into a wing and possibly restore eye development. When *Pc* is knocked down in either mutant background, the eye-to-wing transformation is indeed suppressed and levels of ectopic *Antp* are further elevated (Figure 6D,E). Not only does this result support a cardinal role for Vg-Sd in the eye-to-wing transformation but it also further demonstrates that Antp is not driving this reprogramming event. The reduction of both *vg* and *Pc* (*vg^1^; DE>Pc RNAi*) results in early pupal lethality (Figure 6D’), while the combined reduction of *sd* and *Pc* (*sd^1^; ey>Pc RNAi*) results in pharate lethal adults with ‘tumorous’ head structures (Figure 6E’).

Surprisingly, *vg^1^; DE>Pc RNAi* and *sd^1^; ey>Pc RNAi* discs undergo dramatic hyperplastic growth. This is reminiscent of eye-antennal discs in which other PcG factors have been depleted (Bunker et al., 2015; Classen et al., 2009; Loubiere et al., 2016; Martinez et al., 2009). Both *vg^1^; DE>Pc RNAi* and *sd^1^; ey>Pc RNAi* discs show patterns of tissue folding and organization that indicate it is being specified into several types of imaginal discs. We used immunohistochemistry and RNA-seq to determine if both discs have adopted fates outside of the eye-antennal disc. We first treated the two types of hyperplastic discs with antibodies that distinguish between eye, antennal, wing, leg, and haltere imaginal discs. The spatial distribution of Wg protein is an excellent indicator of wing versus haltere fate. In both wing and haltere discs, *wg* is expressed in a stripe across and around the pouch, however the shape of the pouch is oval in the wing disc and circular in the haltere disc. Differences in the shape of the *wg* expression pattern makes it relatively easy to distinguish these two tissues from each other. The spatial distribution of Wg protein can normally be used to distinguish the antenna from the leg as it is differentially expressed within these discs (Figure 7A). However, since the eye-antennal disc has undergone hyperplastic growth, it is grossly deformed, and *wg* expression alone cannot be reliably used to identify regions of the disc that might have acquired these fates. Therefore, to better distinguish between putative antennal and leg tissue we instead treated hyperplastic discs with antibodies that recognize Dachshund (Dac). Dac protein is distributed in several concentric rings within the leg disc while its spatial expression pattern within the antennal field resembles a signet ring (Figure 7D). The spatial distribution of Wg and Dac proteins within both *vg^1^; DE>Pc RNAi* and *sd^1^; ey>Pc RNAi* eye-antennal discs indicates that a surprising level of heterogeneity exists within the hyperplastic tissue (Figure 7B,C,E,F). An individual disc can simultaneously contain eye, antenna, leg, wing, and haltere tissue (Figure 7G,H).

**Figure 7.**
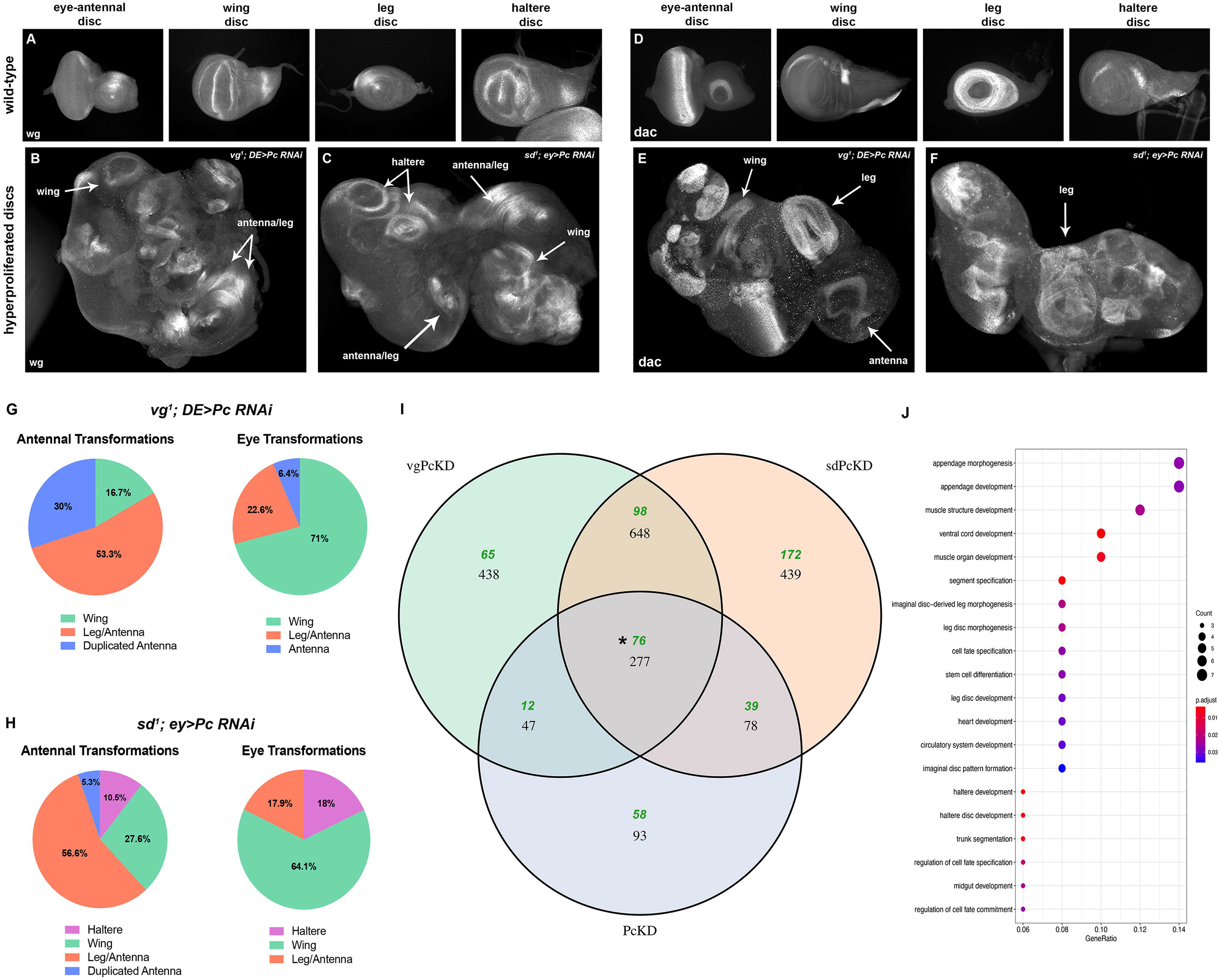
Fate transformations occur in hyperplastic tissue. (A) Wild-type third instar eye-antennal, wing, leg, and haltere imaginal discs stained with anti-Wingless (Wg). (B-C) When *Pc* is lost in *vg^1^* (B) or *sd^1^*(C) mutants, the eye-to-wing transformation is disrupted, and the tissue instead shows evidence of specifying other imaginal fates (arrows). (D) Wild-type third instar eye-antennal, wing, leg, and haltere imaginal discs stained with anti-Dachshund (Dac). (E-F) When *Pc* is lost in *vg^1^* (E) or *sd^1^* (F) mutants, discs specify other imaginal fates (arrows). (G) The loss of *vg* and *Pc* results in high instances of wing specification with additional antennal or leg transformations in the eye and antennal fields. (H) The combined loss of *sd* and *Pc* results in similar novel fate specification, with unique instances of haltere specification in both the eye and antennal fields. (I) Venn diagram comparing the differentially expressed genes of *ey>Pc RNAi* (*PcKD*), *vg^1^; DE>Pc RNAi* (*vgPcKD*), and *sd^1^; ey>Pc RNAi* (*sdPcKD*) eye-antennal discs. Green text represents the number of enriched genes and black represents total number of differentially expressed genes. (J) Enriched Gene Ontology (GO) analysis of overlapping enriched genes between *vgPcKD*, *sdPcKD*, and *PcKD* discs (asterisk in I).

Using bulk RNA-seq we determined the transcriptomes of *vg^1^; ey>Pc RNAi (vgPcKD)* and *sd^1^; ey>Pc RNAi (sdPcKD)* eye-antennal discs and compared these profiles back to those of wild-type and *ey>Pc RNAi (PcKD)* eye-antennal discs (Figure 7I, Table S4). Comparing the lists of differentially expressed genes to each other allowed us to identify potential molecular signatures underlying unique changes in tissue specification and proliferation. *vgPcKD* and *sdPcKD* discs share the differential expression of 648 genes (Figure 7I, Tables S4-6). This is expected as both types of discs show hyperplastic growth and undergo transdetermination events (Figure 7B-H). Interestingly, the percentage of unique fate transformations within the eye and antennal fields differ between *vgPcKD* and *sdPcKD* discs and the emergence of haltere tissue appears only within *sdPcKD* discs (Figure 7B-H). As such, both types of mutant discs also contain unique sets of differentially expressed genes. For example, *vgPcKD* discs are characterized by the differential expression of a unique set of 438 genes while the expression of a completely different set of 439 genes is specifically altered in *sdPcKD* discs (Figure 7I, Tables S4-6). These phenotypic and molecular differences were surprising as disrupting the Vg-Sd complex with either *vg* or *sd* loss-of-function mutants were expected to result in very similar mutant phenotypes and changes in gene expression. The differences that we see might be explained by the fact Sd and Vg are not obligate partners; therefore, mutating each one is likely disrupting different sets of biochemical complexes. We also note that that despite the phenotypic differences between *vgPcKD*, *sdPcKD*, and *PcKD* discs, all three mutant tissues share the differential expression of 277 genes (Figure 7I, Tables S4-6). GO analysis of the elevated genes identified enrichment clusters for imaginal disc development, morphogenesis, and cell fate specification (Figure 7J).

To further infer the trends in gene expression behind the varying disc phenotypes, we categorized each upregulated and downregulated gene based on its primary annotated function (Tables S5,6). Since the molecular and biological function of a significant fraction of these genes have not been elucidated, we looked at the remaining annotated genes for molecular clues that would explain the observed changes in tissue fate and growth. We focused on Hox, selector, and cell proliferation genes since these are most likely to induce the mutant phenotypes seen in our study. The most notable genes are depicted in Figure 8C such as the Hox gene *Antp*, which disrupts eye development (Figure 2), and the wing selector gene *vg,* which transforms it into a wing (Figure 5).

**Figure 8.**
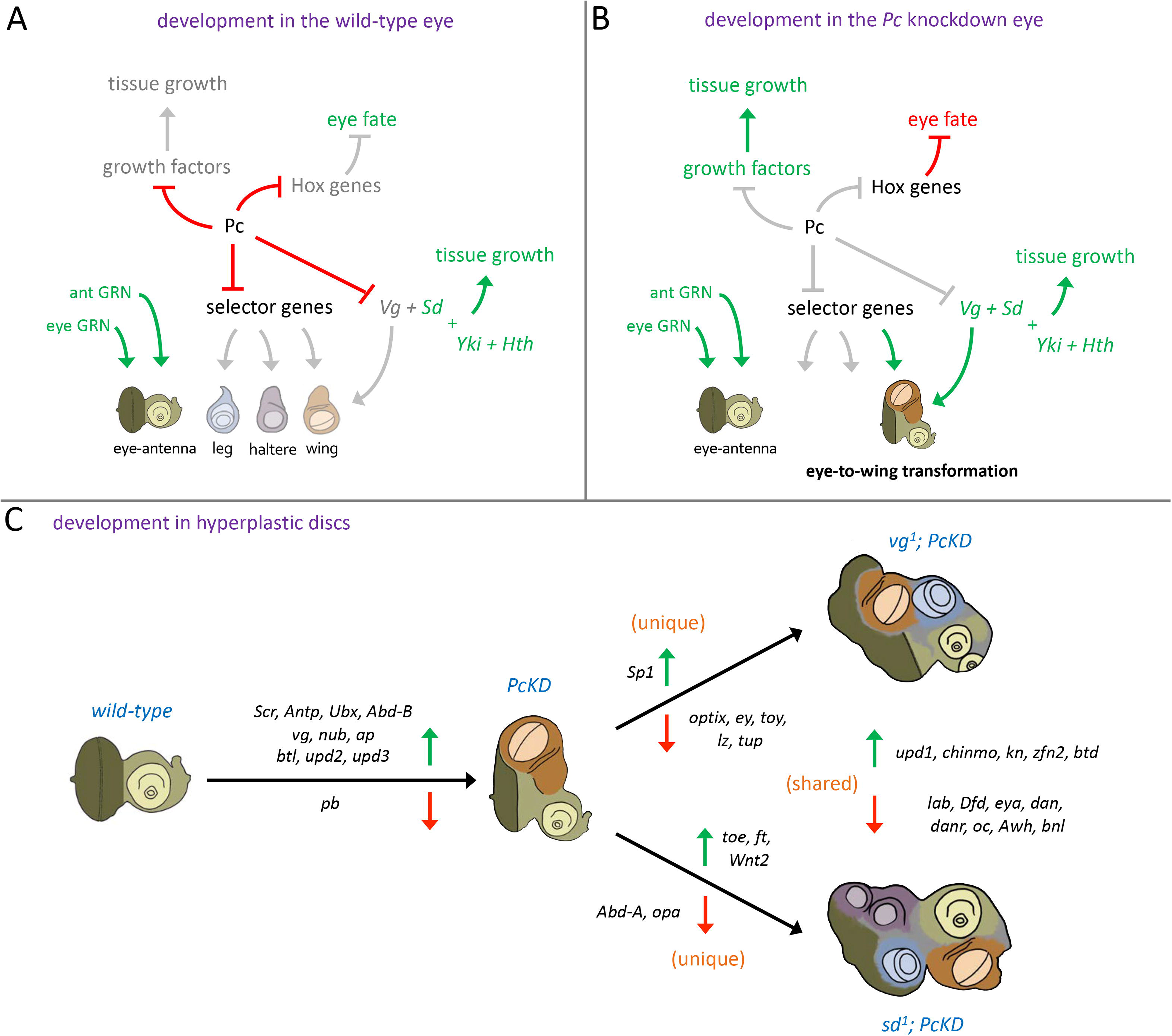
Model for Pc regulation of eye-antennal disc fate. We propose that Pc maintains the fate of the wild-type eye-antennal disc (A) by controlling the expression of genes that encode growth factors, Hox, and selector genes for other imaginal discs. The repression of Vg allows for Sd to form a growth promoting complex with Yki instead of directing wing fate. (B) When *Pc* is knocked down, *vg* expression is activated along with several Hox genes and growth factors. The formation of the Vg-Sd complex transforms the eye into a wing. (C) If the *Pc* knockdown is combined with mutants in *vg* or *sd*, the discs undergo hyperplastic growth which is then organized into other imaginal discs. The immense growth and unique fate specification seen in these discs is reminiscent of intratumor heterogeneity of human tumors and cancers.

The dramatic specification events seen in *vgPcKD* and *sdPcKD* discs is likely caused by a combination of factors. One such factor is the depression of eye and head selector genes (Figure 8C, Tables S5,6) which is known to return it to a multipotent state (Dominguez and Casares, 2005). For example, the loss of RD network members (which we observe in both *vgPcKD* and *sdPcKD* discs) is required for the eye to be transformed into head epidermis or an antenna (Duong et al., 2008; Weasner and Kumar, 2013). Similarly, the loss of *Arrowhead* (*Awh*) (which we also see in both *vgPcKD* and *sdPcKD* discs) allows for the transdetermination of the antenna and head epidermis into an eye (Curtiss and Heilig, 1995; Roignant et al., 2010). Another factor is the ectopic presence of several Hox proteins (Figure 8C, Tables S5,6). Two of these (Antp and Pb) contribute to the inhibition of eye development by physically binding to and inactivating several Pax6 proteins (Benassayag et al., 2003; Plaza et al., 2008; Plaza et al., 2001). The presence of multiple Hox genes in the mutant eye-antennal disc likely contributes to its transformation into other imaginal discs by inhibiting eye development and possibly reverting it to a more multipotent state through the loss or inactivation of RD network factors (7A-H). A final contributing factor disc may be the enrichment of non-ocular selector genes. For example, we have shown that the upregulation of the wing selector gene *vg* transforms the eye into a wing (Figure 5). The enrichment of other wing selector genes such as *nub*, *ap*, and *wg* (Figure 8C, Tables S5,6) are likely needed to sustain wing formation. Similarly, *Sp1* is enriched in *sdPcKD* discs (Figure 8C, Tables S5,6). It serves as a selector gene that promotes leg development and represses wing formation (Ing et al., 2013). In total, the heterogeneity in gene expression likely contributes to the different tissue fates that can be specified only when the Vg-Sd complex is disrupted with either *sd* or *vg* mutants.

Finally, *PcKD* discs show increased tissue growth to accompany the eye-to-wing transformation (Figure 1) (Zhu et al., 2018). Likewise, *vgPcKD* and *sdPcKD* discs are characterized by dramatic hyperplastic growth. Our RNA-seq data indicates that each of these changes in tissue growth are associated with the differential expression of several cell proliferation genes (Figure 8C, Tables S5,6). Of note are the JAK/STAT ligands *upd1*, *upd2*, and *upd3*. The latter two are first upregulated during the transdetermination of an eye into a wing while the other, *upd1*, is enriched solely in the hyperplastic *vgPcKD* and *sdPcKD* discs (Figure 8C, Tables S5,6). The *upd1* and *upd2* genes both contain functional PRE elements and are associated with hyperplasia in imaginal discs (Gonzalez and Busturia, 2009; Nie et al., 2019). As such, these are potentially the main drivers of the increased proliferation that we see in the *PcKD*, *vgPcKD* and *sdPcKD* mutant discs.

## Discussion

For decades disc fragmentation studies, phenotypic analysis of genetic mutants, and over-expression studies have suggested that the eye disc is unusual when compared to other imaginal discs in that it has a limited capacity to have its fate altered. Even when it is redirected towards a different fate it appears to always adopt the fate of a wing (Gehring, 1978; Hadorn, 1968, 1978; Weasner and Kumar, 2022). In this manuscript we provide a mechanism that explains why the eye predominantly transforms into a wing and offer the surprising observation that, under the right circumstances, the eye will adopt a multitude of other imaginal disc fates. Our starting point into this paradigm is the observation that knocking down the epigenetic repressor *Pc* induces the transformation of the eye into a wing (Zhu et al., 2018). In that report, it was proposed that a de-repression of the Hox gene *Antp* (caused by reductions in *Pc*) is responsible for the eye-to-wing transformation. This was a viable hypothesis since (1) *Antp* is expressed broadly within the normal developing wing (Carroll et al., 1995; Paul et al., 2021); (2) reductions in *Antp* expression inhibits proper wing development (Fang et al., 2022; Paul et al., 2021; Struhl, 1981); (3) Pc directly regulates *Antp* expression via functional PRE elements (Ringrose et al., 2003; Zink et al., 1991); (4) *Antp* expression is upregulated in *Pc* knockdown eye discs (Zhu et al., 2018); and (5) the eye-to-wing transformation is observed in a gain-of-function allele of *Antp* (Prince et al., 2008; Scott et al., 1983).

However, despite these observations there are several reasons to doubt the centrality of *Antp* in this process. First, we show in this manuscript that eye-to-wing transformations are not recovered when the expression of other PcG members are reduced even though *Antp* expression is upregulated within the eye field (Figure 1). Second, eye-to-wing transformations are not associated with the vast majority of *Antp* gain-of-function alleles – the only exception being the Cephalathorax (Ctx) allele (Prince et al., 2008; Scott et al., 1983). Instead, gain-of-function *Antp* alleles must be combined with loss-of-function alleles in the Pax6 gene *twin of eyeless* (*toy*) in order to induce the eye-to-wing transformation (Gehring et al., 2009; Papadopoulos et al., 2011). Third, the forced expression of *Antp* on its own within the eye disc is insufficient to induce the eye-to-wing transformation (Figure 2) (Jorgensen and Garber, 1987; Schneuwly et al., 1987a). For this fate change to take place, the over-expression of *Antp* must be accompanied by the hyper-activation of the Notch signaling pathway (Katsuyama et al., 2005; Kurata et al., 2000; Prince et al., 2008). We show that this limitation is not restricted to *Antp* as forced expression of any Hox gene seems to simply disrupt eye development rather than inducing inter-disc fate transformations (Figure 2). Interestingly, over-expression of Hox genes can drive the reprogramming of vertebrate skeletal progenitors into a more uncommitted stem-like state (Leclerc et al., 2023). Since the evidence for *Antp* playing a role in the eye-to-wing transformation is mixed, we set out to identify an alternative explanation for why the eye transforms into a wing when *Pc* levels are reduced.

We compared the transcriptome and epigenome profiles of wild-type eye-antennal discs, wild-type wing discs, and *Pc* knockdown eye-antennal discs to each other and identified several genes that have higher transcript counts and lower H3K27me3 levels throughout the third larval instar (Figures 3,4). Of these, the *vg* gene merited special consideration as it is a known wing selector gene (Kim et al., 1996; Williams et al., 1991), is transcriptionally silent in the wild-type eye disc (Zhu et al., 2018), contains two functional PRE elements (Ahmad and Spens, 2019; Okulski et al., 2011; Ringrose et al., 2003), and induces an eye-to-wing transformation when ectopically expressed within the eye field (Figure 5) (Kim et al., 1996; Simmonds et al., 1998). We note here that PcG-mediated regulation of additional genes is important for maintaining the fate of the eye because the expression of H3K27M histone proteins (which cannot be methylated) interferes and blocks Vg-mediated eye-to-wing transformation (Ahmad and Henikoff, 2021).

We then investigated the mechanism by which ectopic Vg alters the fate of the eye. Vg is a transcriptional activator (MacKay et al., 2003) that binds to the Sd DNA binding protein and promotes wing formation (Halder et al., 1998; Simmonds et al., 1998). The binding of Vg to Sd appears to alter the enhancer specificity of the Sd protein and forces it to bind to only wing specific target genes (Garg and Bell, 2010; Halder and Carroll, 2001). This implies that in tissues such as the eye, where Vg is absent (Figure 5) (Zhu et al., 2018), Sd will bind to a distinct set of genes that are not involved in wing formation. Indeed, several Vestigial independent roles have been identified for Sd in diverse tissues such as the larval eye and leg, as well as the adult central nervous system (Garg et al., 2007). Within the eye, the Yorkie (Yki) transcriptional activator binds to Sd (Yki-Sd) to relieve default repression of growth promoting genes and to promote re-entry of quiescent cells into the cell cycle during tissue regeneration (Koontz et al., 2013; Meserve and Duronio, 2015; Wu et al., 2008). Our model is that ectopic activation of Vg within the eye (due to reductions in *Pc* levels) allows for Vg to bind Sd and redirect it away from cell proliferation genes and onto wing fate promoting genes instead (Figure 8A,B). A similar model has been proposed for the wing itself where the formation of Vg-Sd complexes within the pouch indirectly reduces tissue growth by titrating Sd away from Yki (Koontz et al., 2013). Our model is supported by the fact that reducing *sd* expression prevents the Vg-mediated eye-to-wing transformation (Figure 6).

Our findings suggest that Pc mediates the eye vs wing decision by selectively repressing *vg* within the developing eye primordium while allowing its expression within the developing wing pouch. While not tested here, our findings also provide a potential mechanism by which the eye is transformed into a wing under other experimental conditions. In each instance, be it regenerating cells of a disc fragment or the endogenous eye in which selector genes are reduced or over-expressed, one possible explanation for why the eye is transformed into a wing involves the de-repression of *vg* within the eye disc. How might *vg* expression be activated in the absence of Pc repression within the eye field? Several studies have shown that, in the developing wing, *vg* expression is activated by the Notch and Wingless signaling pathways (Bernard et al., 2009; Couso et al., 1995; Djiane et al., 2014; Koelzer and Klein, 2006; Neumann and Cohen, 1996; Zecca and Struhl, 2007a, b). Similarly, *vg* expression is activated within the eye when *Antp* and the Notch pathway are simultaneously activated but not when *Antp* is ectopically expressed alone (Katsuyama et al., 2005; Kurata et al., 2000). Based on these prior studies, we propose that the Notch and Wingless pathways, which play important roles in the developing eye (Baker, 2007; Silver and Rebay, 2005; Tio et al., 1996; Voas and Rebay, 2004), could activate *vg* expression in the absence of *Pc* mediated repression. Under these conditions, the formation of the Vg-Sd complex would shift the balance away from eye development (Yki-Sd) and towards wing formation (Vg-Sd).

To further test our model, we used *vg* and *sd* loss-of-function mutants to disrupt the Vg-Sd complex while simultaneously reducing *Pc* expression levels. Our expectation was that the eye-to-wing transformation would be simply blocked, and eye development might even be restored. While the percentage of eye-to-wing transformations was reduced, we surprisingly observed that the mutant eye-antennal discs underwent dramatic levels of hyperplastic growth (Figure 6). This tumor-like growth is reminiscent of discs in which other PcG members have been depleted (Classen et al., 2009; Medina et al., 2021) or when epithelial polarity is lost within the eye-antennal disc (Bunker et al., 2015; Enomoto et al., 2021; Pagliarini and Xu, 2003; Wu et al., 2010). We have further made the unexpected discovery that the *vgPcKD* and *sdPcKD* hyperplastic discs do not just consist of amorphous tissue but instead are regionally specified as other imaginal discs (Figure 7). Our bulk RNA-seq analysis of these mutant eye-antennal discs suggest that once the primary push towards a wing fate is eliminated (through disruption of the Vg-Sd complex), cell proliferation genes likely drive hyperplastic growth while changes in Hox and selector genes disrupt the pre-programmed fate and force the tissue to transdetermine into several new imaginal disc fates, respectively. (Figure 7, Table S5,6). Interestingly, the JAK/STAT ligand, Upd1, whose levels are elevated in both *vgPcKD* and *sdPcKD* hyperplastic discs, has recently been associated with intratumor heterogeneity in *Drosophila* follicle cell tumors (Chatterjee et al., 2023). The diversity in tissue fates that we observe in the *vgPcKD* and *sdPcKD* discs is also reminiscent of intratumor heterogeneity that is seen in a wide range of human tumors (Ju, 2021; Li et al., 2022; McGranahan and Swanton, 2017). As such, the induction of hyperplastic imaginal discs could serve as tractable model systems for elucidating the genetic and molecular mechanisms that underly tumor heterogeneity.

Based on the bulk RNA-seq results throughout our study, we have in Figure 8C summarized potentially important changes in gene expression that could underlie the alterations in tissue fate we observe when we knockdown *Pc* alone or in combination with a disruption of the Vg-Sd complex. In this model we have focused on changes to the expression levels of Hox genes, tissue selector genes, and tissue proliferation genes. These classes of genes are reasonable to consider since (1) altering Hox and eye selector gene expression inhibits eye development; (2) raising transcription levels of wing selector genes is sufficient to transform the eye into a wing; (3) altering expression levels of cell proliferation genes is associated with elevated tissue growth and tumorigenesis; and (4) the forced expression of non-ocular selector genes is known to induce homeotic transformations. While the differential expression of the listed genes alone is unlikely to account for the dramatic changes in tissue fate and growth control, they provide insight into the mechanisms by which *Pc* expression maintains the fate of the eye-antennal disc.

### Materials and Methods Genetics

The following fly stocks for experiments in the main text were obtained from the Bloomington *Drosophila* Stock Center (BDSC), Indiana University, Bloomington, USA: *ey-GAL4* 8221, *UAS-Pc RNAi* 33964, *UAS-Scr* 7302, *UAS-Antp* 7301, *UAS-Ubx* 911, *UAS-Abd-B* 913, *UAS-lab* 7300, *UAS-pb* 7298, *UAS-abd-A* 912, *UAS-Dfd* 7299, *UAS-pho RNAi* 42926, *UAS-Sfmbt RNAi* 28677, *UAS-calypso RNAi* 56888, *UAS-Sce RNAi* 67924, *UAS-Scm RNAi* 55278, *UAS-vg* 37296, *vg^1^* 432, *sd^1^* 1027, *UAS-sd RNAi* 29352, *c311-GAL4* 5937, and *DE-GAL4* (Georg Halder, Katholic University, Leuven, Belgium). Supplemental screening stocks are noted in *Supplemental Table 1*. All crosses were carried out at 25°C. For scoring of all eye-antennal discs and adult phenotypes, n=30.

### Immunofluorescence and imaging

The following antibodies were used: rat anti-Elav (Developmental Studies Hybridoma Bank [DSHB] #7E8A10, 1:100), mouse anti-Antp (DSHB #8C11, 1:100), mouse anti-Wg (DSHB #4D4, 1:800), mouse anti-Dac (DSHB #2-3, 1:5), rabbit anti-Vg (K. Guss, Dickinson College, Carlisle, PA, 1:100), FITC-conjugated donkey anti-rabbit (Jackson #711-095-152, 1:100), FITC-conjugated donkey anti-rat (Jackson #712-095-153, 1:100), Cy3-conjugated donkey anti-mouse (Jackson #715-165-151), Alexa Fluor 647-conjugated phalloidin (Thermo #A22287, 1:20), and Rhodamine phalloidin (Thermo #R415, 1:100). *Drosophila* eye-antennal imaginal discs were prepared as previously described (Spratford and Kumar, 2014). Fluorescent images were taken using a Zeiss Axioplan II compound microscope and processed by Fiji/ImageJ (Schindelin et al., 2012) and Adobe Photoshop software. Adult flies were imaged with either a Zeiss Discovery V12 or a Leica M205FA stereo microscope.

### Molecular biology

#### RNA-seq

Samples were collected from eye-antennal or wing discs throughout third instar development (84hr-144hr after egg lay). 50 eye-antennal imaginal discs were dissected from *ey>Pc RNAi* or *w^1118^* larvae, 20 wing discs from *w^1118^* larvae, or 25 eye-antennal discs from *vg^1^; DE>Pc RNAi* or *sd^1^; ey>Pc RNAi* larvae were dissected as previously described (Spratford and Kumar, 2014). Total RNA was extracted with the RNeasy Plus Mini Kit (Qiagen #74134) per kit protocol. 0.3ug RNA was used to generate a polyA-strand specific library with the Illumina TruSeq Stranded mRNA library prep kit. Libraries were prepared and sequenced by the Center for Genomics and Bioinformatics (CGB, Indiana University, Bloomington, USA).

#### CUT&RUN

Chromatin was extracted and purified from ten third-instar eye-antennal or wing imaginal discs as previously described (Weasner et al. 2023). The following antibodies were used: rabbit anti-trimethyl-histone H3 (Lys27) (C36B11) (Cell Signaling #9733, 1:51), rabbit anti-Ubiquityl-Histone H2A (Lys119) (D27C4) (Cell Signaling #8240, 1:51), and Normal Rabbit IgG (Cell Signaling #2729, 1:51). Cleaved chromatin fragments purified with the MinElute PCR Purification Kit (Qiagen #28004) per kit protocol.

### Computational Analysis

#### RNA-seq

Read quality was assessed with *fastqc* (Andrews, 2010) (v0.11.9). Genome indices were generated via *STAR* (Dobin et al., 2013) and were aligned to the dm6 *Drosophila* genome (with the additional setting *--genomeSAindexNbases* 13 for *Drosophila*). Aligned reads were counted using *Subread* (Liao et al., 2019) function f*eatureCounts* with the settings *-F GTF -t exon -g gene_id -- minOverlap 10 --largestOverlap --primary -s 2 -T*. bigWig files of RNA-seq counts were generated with *deepTools* (Ramirez et al., 2016) (v3.5.1) function *bamCoverage* with the settings *--normalizeUsing CPM -bs 1 --smoothLength 25 --filterRNAstrand [forward or reverse] -p 16*. Downstream analysis was performed in Rstudio (v4.2.1). Differential expression analysis was performed with *DESeq2* LO and heatmaps were generated with *ComplexHeatmap* (Gu et al., 2016). Differentially expressed transcripts were filtered with *dplyr* package (Wickham et al., 2022) to select for genes ‘enriched in WD’ (in ead_vs_wd_change) and ‘enriched in absence of Pc’ (in pckd_vs_ead_change). Candidate overlap between each time point were then compared with *VennDiagram* (Chen and Boutros, 2011). Gene ontology analysis was performed with *clusterProfiler* (Yu et al., 2012) *eGO* function (settings: *OrgDb = “org.Dm.eg.db”, ont = “BP”, pAdjustMethod = “fdr”, keyType = “SYMBOL”*). Additional scripts for downstream analysis can be found in supplemental information.

#### CUT&RUN

Read quality was assessed with fastqc (Andrews, 2010) (v0.11.9). Genome indices were generated via *Bowtie2* (Langmead et al., 2009) and were aligned to the *Drosophila* dm6 genome and *E. coli* EB1 genomes (with the settings *min_fragment = 10 max_fragment = 700*). The quality and consistency of raw aligned reads were assessed with deeptools (v3.5.1) function *multiBamSummary* and *plotCorrelation* (with the settings *--whatToPlot heatmap --corMethod spearman --plotNumbers*). The number of *E. coli* aligned reads were accessed with *samtools* (Danecek et al., 2021) (v1.15.1) command *view -c -F 4* and used to generate an appropriate scale factor with the calculation: 10,000/[# *E. coli* aligned reads]. Normalized bigWig and bedgraph files of *Drosophila* CUT&RUN reads were generated with *deepTools* (Ramirez et al., 2016) (v3.9/3.5.1) function *bamCoverage* with the corresponding *-- scaleFactor and --outFileFormat [bigwig or bedgraph]* setting. Initial visualization of bigWig reads was carried out in Integrative Genomics Viewer (v2.8.0); figures of chromatin tracks were generated using *Gviz* (Hahne and Ivanek, 2016) in Rstudio (v4.2.1). Peaks of normalized bedgraph tracks were called with *SEACR* (Meers et al., 2019) web interface (“norm” settings). *deepTools* functions were used to generate all downstream analysis: Peak matrices were generated with *computeMatrix reference-point –referencePoint center –missingDataAsZero -a 2000 -b 2000*. Heatmap of CUT&RUN peaks was generated from matrix files with *plotHeatmap --matrixFile --whatToShow ‘plot, heatmap, and colorbar’*. Additional scripts for downstream analysis can be found in supplemental information.

## Acknowledgements

The authors are grateful to Kirsten Guss for the anti-Vg antibody, to the Developmental Studies Hybridoma Bank (DSHB) for other primary antibodies listed within the Key Resource Table, to Georg Halder for the DE-GAL4 fly strain, to the Bloomington Drosophila Stock Center for all remaining fly strains listed within the Key Resource Table, to both Gabe Zentner and Robert Policastro for technical support with bioinformatics, to the Light Microscopy Imaging Center (LMIC) for use of the Leica M205FA stereo microscope, and to the Center for Genomics and Bioinformatics (CGB) for the generation of sequencing libraries and next-generation sequencing.

## Competing Interests

The authors declare no competing interests.

## Author Contributions

Conceptualization, H.E.B. and J.P.K.; Data curation, H.E.B.; Formal analysis H.E.B.; Funding acquisition, J.P.K.; Investigation, H.E.B, B.P.W., and B.M.W.; Methodology, H.E.B. and B.P.W.; Software, H.E.B.; Visualization, H.E.B. and J.P.K.; Writing – original draft, H.E.B. and J.P.K.; Writing – review and editing, H.E.B., B.M.W., and J.P.K.

## Funding

This work is supported by the Indiana University Robert W. Briggs Summer Fellowship in Developmental Biology to Haley E. Brown and a research grant from the National Eye Institute (R01 EY030847) to Justin

P. Kumar.

## Data availability

Scripts used for the initial RNA-seq (https://github.com/rpolicastro/RNAseq) and CUT&RUN (https://github.com/gzentner/ChIPseq) pipelines can be found on Github. Additional scripts for downstream analysis can be found in supplemental information. Raw data files can be accessed at SRA accession number PRJNA949271. File are also available upon request.

## Notes

### Competing Interest Statement

The authors have declared no competing interest.

